# Weight Transfer in the Reinforcement Learning Model of Songbird Acquisition

**DOI:** 10.1101/2024.12.30.628217

**Authors:** Khue Tran, Alexei Koulakov

## Abstract

Song acquisition behavior observed in the songbird system provides a notable example of learning through trial-and-error which parallels human speech acquisition. Studying songbird vocal learning can offer insights into mechanisms underlying human language. We present a computational model of song learning that integrates reinforcement learning (RL) and Hebbian learning and agrees with known songbird circuitry. The song circuit outputs activity from nucleus RA, which receives two primary inputs: timing information from area HVC and stochastic activity from nucleus LMAN. Additionally, song learning relies on Area X, a basal ganglia area that receives dopaminergic inputs from VTA. In our model, song is first acquired in the HVC-to-Area X connectivity, employing an RL mechanism that involves node perturbation. This information is then consolidated into HVC-to-RA synapses through a Hebbian mechanism. The transfer of weights from Area X to RA takes place via the thalamus, utilizing a specific form of spike-timing-dependent plasticity (STDP). Thus, we present a computational model grounded in songbird circuitry in which the optimal policy is initially guided by RL and subsequently transferred to another circuit through Hebbian plasticity.

## I. Introduction

Juvenile songbirds faithfully learn to reproduce a tutor song via repeated vocal babbling, a process that illustrates robust learning by experimentation [1] and mirrors human speech acquisition [2]. Comparative studies have analyzed similarities in gene expression involved in vocal learning of songbirds and humans [2]. These observations argue for the convergent evolution of song learning and speech acquisition circuits. The high degree of homology shared between the songbird and human systems motivates models of song learning, which provide further insights into human language models. We want to study the network underlying song acquisition behavior as a model system for sensorimotor learning in the brain, which can also inform the development of machine learning algorithms for human language.

As summarized in Figure 1, experimental studies have determined two prominent pathways for songbird vocal acquisition: song production and song learning. The song production pathway includes the projection from HVC, a region with stereotyped activity patterns that provide timing information, to RA, a premotor area that ultimately drives downstream motor outputs [3]. The song learning pathway follows the HVC connectivity to Area X, a basal ganglia region that crucially receives dopaminergic inputs from VTA [3][4]. The two pathways interact via nucleus LMAN, which synapses to RA and receives inputs from Area X through the thalamus [3]. LMAN neurons provide stochastic noise to the network, which is necessary for juvenile birds to generate variability for practice songs [3][5].

**Fig. 1.**
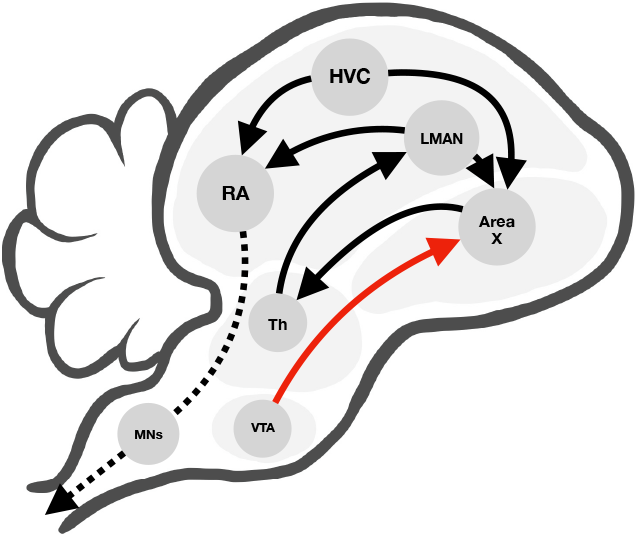
Schematic of songbird anatomy with diverging pathways for song production and learning (black arrows). The song production pathway follows the projection from HVC to RA, then from RA to motor neurons which generate behavior. The song learning pathway follows the connectivity from HVC to Area X, which forms a feedback loop with LMAN and connects to the production pathway. Area X also receives direct inputs from VTA (red arrow).

**Fig. 2.**
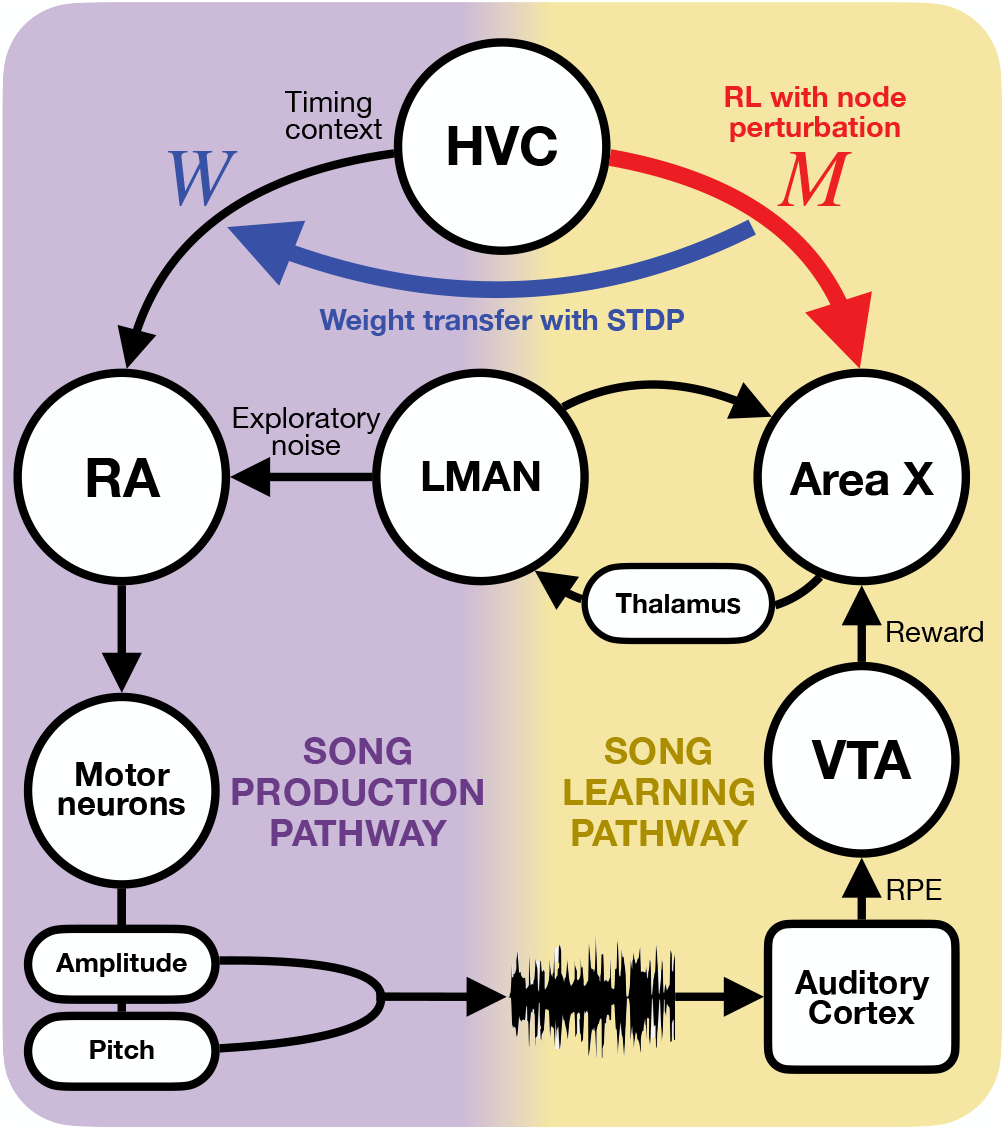
The architecture of our model includes the song production and song learning pathways. Learning at HVC-to-Area X synapses (red), described by the matrix *M* , is enabled with timing context from HVC, reward from VTA, and exploratory noise form LMAN. Weight transfer (blue) from *M* to HVC-to-RA synapses, *W* , is facilitated by an form of unsupervised Hebbian learning, utilizing an STDP kernel that combines the presynaptic input from HVC and postsynaptic response of RA.

**Fig. 3.**
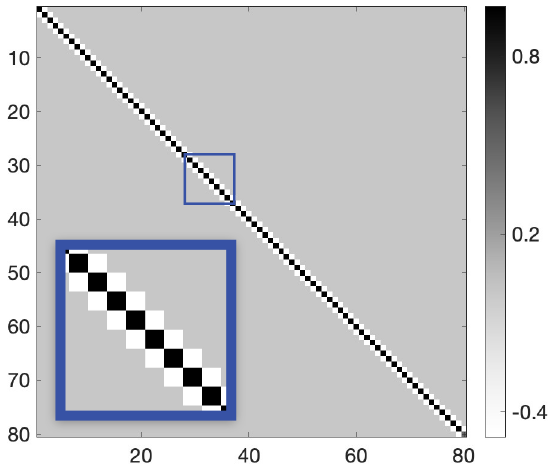
STDP kernel *K* for weight transfer

Previously, Doya and Sejnowski demonstrated that a model following songbird circuitry can be trained to learn a tutor song using weight perturbation. The model was a three-layer feed-forward network that resembles the song production pathway. Reinforcement learning (RL) at HVC-to-RA synapses was enabled with exploratory noise from LMAN and reinforcement signals from VTA, both projecting to RA [6][7]. In 2007, Fiete et al. revised the model and showed that node perturbation could also be used to train a similar network [8].

There remains a gap between existing models and the observed circuitry where the primary locus of RL is suggested to be at Area X in the song learning pathway [3][4][9]. It is unknown how the learned information is consolidated into the song production pathway. We present a computational model for learning that includes the acquisition of song sequence in the song learning pathway with RL and then transferred to the song production pathway via spike-timing-dependent plasticity (STDP).

## II. Model Description

Our model follows the hypothesis that there is a critical period of learning where HVC-to-Area X weights are updated trough an RL-based mechanism. Subsequently, weights are transferred to HVC-to-RA synapses where this information is crystallized into adulthood.

HVC activity can be represented by an *N* ×*T* matrix **H** = [*h*_1_, …, *h*_*T*_ ], where *N* is the number of HVC neurons and *T* is the total number of time steps. Each column in this matrix defines the response of the HVC population at a given time point. We will assume the following form for the response matrix, implying that each HVC neuron produces a transient peak of activity at a specific time within the song:

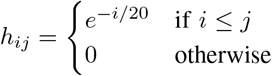

We will represent Area X activity in the *M* × *T* matrix **X** = [*x*_1_, …, *x*_*T*_ ] with each column, *x*_*t*_, representing the firing rate of *M* neurons at time step *t*. Given the feedforward projection from HVC to Area X defined by the weight matrix **M** ∈ ℝ^*M* ×*N*^ , one can write the following equation describing activity of Area X neurons:

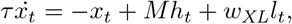

where *τ* is the synaptic time constant and *w*_*XL*_ is the scalar describing the connectivity strength between LMAN and Area X. Area X receives inputs from region LMAN, whose activity is defined by **L** = [*l*_1_, …, *l*_*T*_ ], a *M* ×*T* matrix. LMAN-to-Area X and Area X-to-LMAN neurons form a feedback loop which is organized tonotopically with scalar weights *w*_*XL*_ = 0.3 and *w*_*LX*_ = 1, respectively. In addition to the feedback, LMAN neurons receive inputs that are randomly sampled at every time step from the normal distribution,

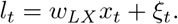

Here, *ξ*_*t*_ ∼ 𝒩 (0, *σ*_LMAN_). In our simulations, we used *σ*_LMAN_ = 0.01.

The responses of RA neurons **R** = [*r*_1_, …, *r*_*T*_ ], an *M* × *T* matrix, are affected by projections from LMAN and direct inputs from HVC through weight matrix **W** ∈ ℝ^*M* ×*N*^ . RA then projects to the population of output motor neurons 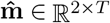, controlling the sound pitch and amplitude, via weight matrix **A** ∈ ℝ^2×*M*^ , which is constant,

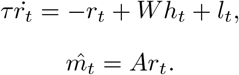

Every time step, an eligibility trace is computed between coactive LMAN and HVC neurons: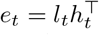. At the end of each iteration, motor outputs are evaluated against tutor signals **m** ∈ ℝ^2×*T*^ to compute the reward 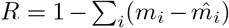 and to update weights.

HVC-to-Area X weights, **M**, are updated with a RL learning rule based on node perturbation with reinforcement from the reward and exploratory noise from LMAN activity [6][8]:

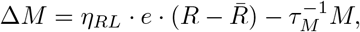

where *η*_*RL*_ is the RL learning rate and *τ*_*M*_ the weight decay time constant.

HVC-to-RA weights, **W**, are learned with an unsupervised STDP learning rule that relies on kernel *K*, the presynaptic activity of HVC, and the postsynaptic activity of RA:

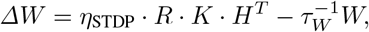

where *η*_STDP_ is the STDP learning rate and *τ*_*W*_ the weight decay time constant.

The kernel, **K** ∈ ℝ^*T* ×*T*^ , is calculated to ensure an exact copying from *M* to *W* ,

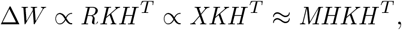

where we solve for *K* by enforcing *HKH* ^*T*^ = *I* , which leads to *W* ∝ *M* thus implementing weight transfer from *M* to *W* .

## III. Results

We trained the model to learn a template song of fixed amplitudes and frequencies (Fig. 4). Training results show that over iterations, the correlations between HVC-to-RA and HVC-to-Area X weights and reward saturate at 1 (Fig. 5). The network was simulated with 160 HVC neurons and 200 RA, LMAN, and Area X neurons. The template song lasted for 80 time steps and weights were updated at the end of each training iteration. After 20,000 iterations, the final song was evaluated with LMAN-to-RA inputs removed. Both trainable weight matrices, *M* and *W* , were randomly initialized and matrix *A* was fixed according to parameters described previously [8]. Training parameters included *η*_*RL*_ = 10^−3^, *η*_*ST DP*_ = 10^−4^, *τ* = 10, *τ*_*M*_ = 10^6^, *τ*_*W*_ = 10^5^.

**Fig. 4.**
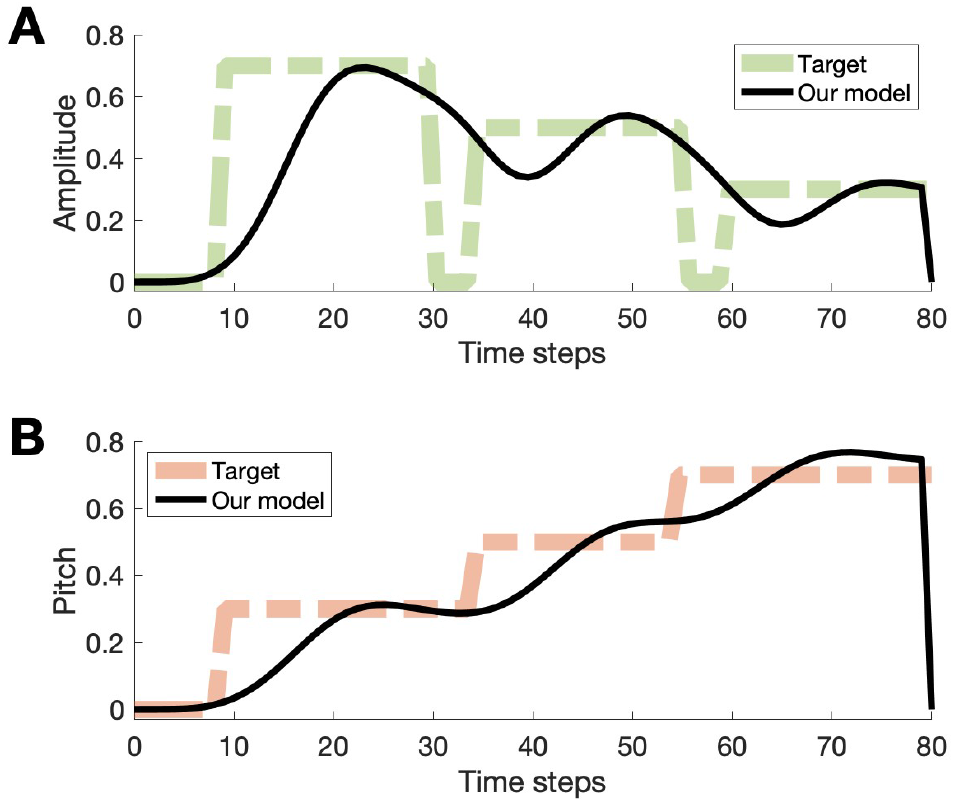
Results from model output. After training the model, LMAN inputs to RA were removed to generate two output traces. A) Target amplitude used to train the model shown in green and the black trace is the output of our model. B) The target pitch trace shown in orange and the result from our model in black.

**Fig. 5.**
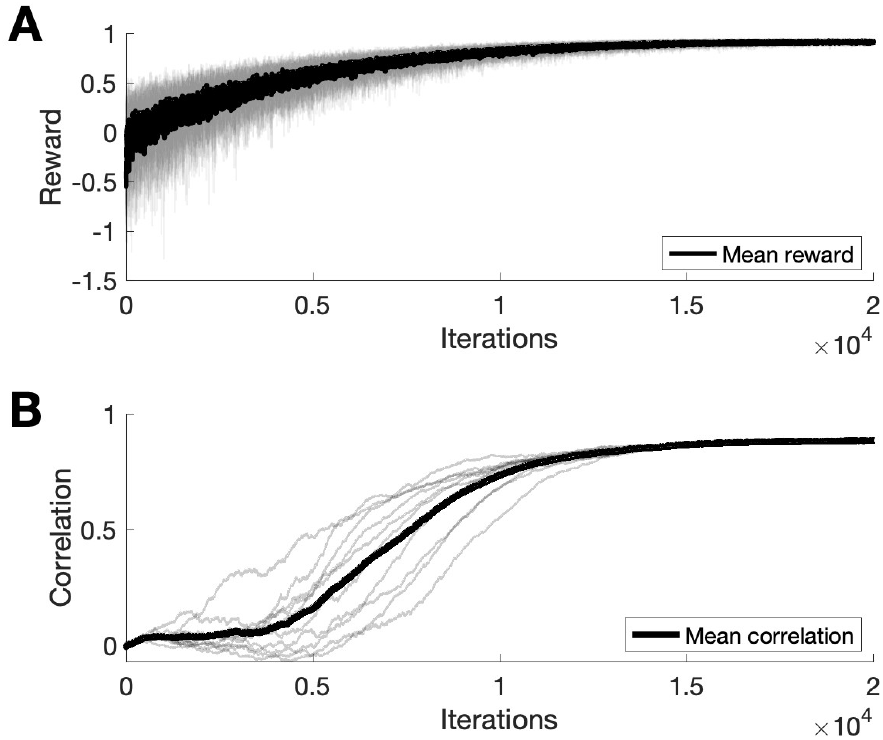
Results from training. A) Reward evaluated from final song increases across training iterations. Black trace shows the mean reward across 10 different runs, individual reward traces are in grey. B) Correlations between HVC-to-RA and HVC-to-Area X weights increase over training iterations, indicating successful weight transfer. Black trace shows the mean correlation across 10 different runs, individual correlations are in grey.

To show that our model of song acquisition requires both node perturbation-based RL in Area X and STDP-facilitated weight transfer to RA, we performed a series of ablation experiments with modifications to the model and compared final reward values after training (Fig. 6). We first confirmed that the model described by Doya and Sejnowski and Fiete et al. achieved baseline performance [6], [8]. By disentangling song production and learning pathway with separate learning mechanisms, our model performs worse in comparison by a small amount. However, with Area X containing the locus of RL, our model performance reflects experimental findings that suggest learning in Area X is required for song acquisition [9].

**Fig. 6.**
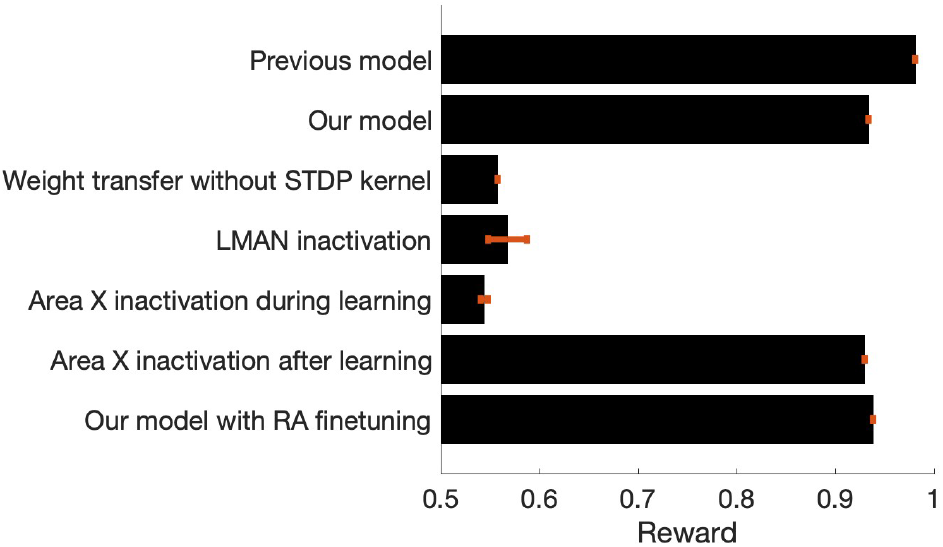
Ablation experiments show both RL and Hebbian plasticity necessary for song acquisition and consolidation. Reward values shown are from the final produced song without noise inputs, bar plot summarizes the mean reward across 10 different runs for each model and SEM shown in red, with maximum reward of 1 indicating perfect learning.

When we disable RL updates to HVC-to-Area X weights early in the training process, a deficit comparable to inactivating of Area X for juvenile songbirds, learning is impaired. But if we limit RL updates to the first 5,000 iterations and then remove the connection from Area X to RA though the thalamus, we observe that song production is unaffected since the weight transfer has already consolidated information into HVC-to-RA synapses by that time point.

In our model, the STDP mechanism is necessary for weight transfer, since removing the kernel impairs final performance. Additionally, if LMAN, the source of exploratory noise, is silenced, the model also cannot learn, which is consistent with experimental findings [3][5].

Lastly, we trained the model to produce a more realistic song from two control signals, analogous to pitch and amplitude, based on the established model for syrinx control [10]. Training parameters to learn this song included 2200 HVC neurons and 300 RA, LMAN, and Area X neurons, *η*_*RL*_ = 10^−3^, *η*_*ST DP*_ = 10^−3^. The model was trained with *T* = 1100 and the outputs were interpolated with time steps of 10^−5^, corresponding to a song length of 1.1 ms. The synthetic song was produced by integrating the model as described by Laje et al. with the two control signals as parameters. We found that our model can learn the simple realistic song (Fig. 7) with somewhat better accuracy than a synthetic pattern containing square pulses (Fig. 4). We suggest that learning temporal sequences via RL with subsequent consolidation in the song production pathway can be accomplished for realistic sequences.

**Fig. 7.**
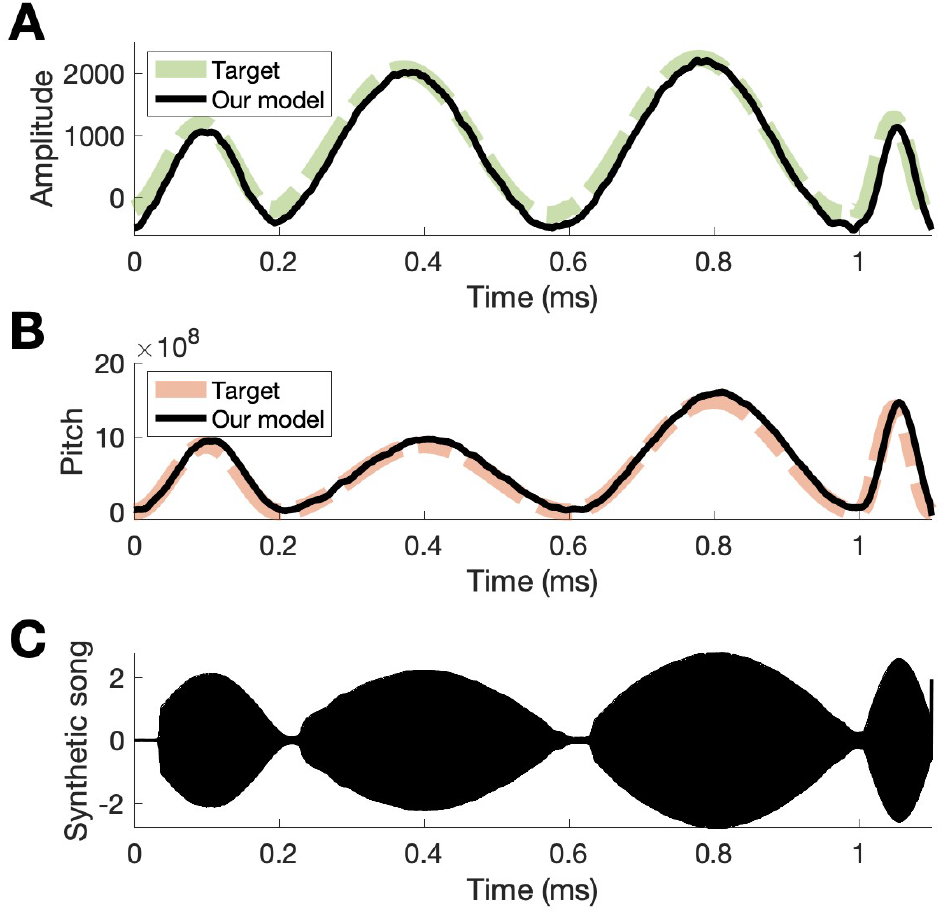
Results from training the model to learn a realistic song. (A), (B) Two parameters learned by the model that correspond the pitch and amplitude of a song. (C). The synthetic song constructed from the two control parameters based on [10].

## IV. Discussion

We have developed a computational model of birdsong vocal learning that combines RL and Hebbian plasticity in the context of known circuitry. In the model, the song learning pathway, specifically the HVC-to-Area X connectivity, initially guides the policy to generate an accurate song with reinforcement from VTA and exploratory noise from LMAN. The song sequence is further consolidated into the song production pathway (HVC-to-RA synapses) via the connection through LMAN. The consolidation of information from Area X to HVC-to-RA synapses relies on Hebbian plasticity with an STDP kernel that facilitates weight transfer. We showed that the model can be used to learn a simple song effectively and weights from the song learning pathway can be consolidated into the production pathway.

The ablation studies confirm that both RL and Hebbian plasticity are essential to the model. The results from our model showed that while RL in Area X is crucial for initial learning, it is no longer essential after the song has been crystallized. This result agrees with the experimental observation of a critical period of learning for juvenile songbirds. With song learning as an example, the model presents a biologically inspired motif of learning and memory consolidation of temporal sequences in the brain.

## V. Acknowledgments

We thank Jesse Goldberg, Drew Schreiner, and Vikram Gadagkar for numerous helpful discussions. This work was supported by NIH BRAIN Initiative grant U19NS112953.

## References

[1] M. S. Fee and C. Scharff, “The songbird as a model for the generation and learning of complex sequential behaviors,” Ilar j, vol. 51, no. 4, pp. 362–77, 2010. DOI: 10.1093/ilar.51.4.362.

[2] M. H. Davenport and E. D. Jarvis, “Birdsong neuroscience and the evolutionary substrates of learned vocalization,” Trends Neurosci, vol. 46, no. 2, pp. 97–99, 2023. DOI: 10.1016/j.tins.2022.11.005.

[3] R. Chen and J. H. Goldberg, “Actor-critic reinforcement learning in the songbird,” Curr Opin Neurobiol, vol. 65, pp. 1–9, 2020. DOI: 10.1016/j.conb.2020.08.005.

[4] M. Fee and J. Goldberg, “A hypothesis for basal ganglia-dependent reinforcement learning in the songbird,” Neuroscience, vol. 198, pp. 152–170, 2011, Function and Dysfunction of the Basal Ganglia, ISSN: 0306-4522. DOI: 10.1016/j.neuroscience.2011.09.069.

[5] T. L. Warren, E. C. Tumer, J. D. Charlesworth, and M. S. Brainard, “Mechanisms and time course of vocal learning and consolidation in the adult songbird,” J Neurophysiol, vol. 106, no. 4, pp. 1806–1821, 2011. DOI: 10.1152/jn.00311.2011.

[6] K. Doya and T. J. Sejnowski, “A novel reinforcement model of birdsong vocalization learning,” in Neural Information Processing Systems, 1994.

[7] V. Gadagkar, P. A. Puzerey, R. Chen, E. Baird-Daniel, A. R. Farhang, and J. H. Goldberg, “Dopamine neurons encode performance error in singing birds,” Science, vol. 354, no. 6317, pp. 1278–1282, 2016. DOI: doi : 10.1126/science.aah6837. [Online]. Available: https://www.science.org/doi/abs/10.1126/science.aah6837.

[8] I. R. Fiete, M. S. Fee, and H. S. Seung, “Model of birdsong learning based on gradient estimation by dynamic perturbation of neural conductances,” Journal of Neurophysiology, vol. 98, no. 4, pp. 2038–2057, 2007. DOI: 10.1152/jn.01311.2006.

[9] C. Scharff and F. Nottebohm, “A comparative study of the behavioral deficits following lesions of various parts of the zebra finch song system: Implications for vocal learning,” J Neurosci, vol. 11, no. 9, pp. 2896–913, 1991. DOI: 10.1523/jneurosci.11-09-02896.1991.

[10] R. Laje, T. J. Gardner, and G. B. Mindlin, “Neuromuscular control of vocalizations in birdsong: A model,” Phys Rev E Stat Nonlin Soft Matter Phys, vol. 65, no. 5 Pt 1, p. 051 921, 2002. DOI: 10.1103/PhysRevE.65.051921.

